# Female meiosis II and pronuclear fusion require Bicaudal-D

**DOI:** 10.1101/2020.12.22.423980

**Authors:** Paula Vazquez-Pianzola, Dirk Beuchle, Gabriella Saro, Greco Hernández, Giovanna Maldonado, Dominique Brunßen, Peter Meister, Beat Suter

## Abstract

*Drosophila* Clathrin heavy chain (Chc) is transported by the dynein/dynactin microtubule motor through its interaction with the adaptor protein Bicaudal-D (BicD). Here we show that *Drosophila* BicD and Chc localize to centrosomes and spindles during mitosis and to the tandem spindles during female meiosis II. Reducing the activity of BicD::GFP specifically in freshly laid eggs revealed that BicD is essential for the production of normal female meiosis II products and for pronuclear fusion. Chc interacts with BicD and D-TACC, and BicD is needed to correctly localize the microtubule-stabilizing factors D-TACC, clathrin, and Msps to the meiosis II spindles, suggesting that BicD acts by localizing these proteins. In unfertilized eggs, reduced BicD levels cause the female meiotic products to re-enter the cell cycle. As BicD is required to localize the spindle assembly checkpoint (SAC) components Mad2 and BubR1 to the female meiotic products, it appears that BicD functions to localize them to control metaphase arrest of polar bodies. Finally, *Drosophila* and *C. elegans* orthologs of *BicD* and *tacc* are also needed for pronuclear fusion.

## INTRODUCTION

Encoded by a single gene, the *Drosophila* BicD protein is part of a family of evolutionarily conserved dynein adaptors responsible for the transport of different cargoes along microtubules (Vazquez-Pianzola et al., 2016; Hoogenraad and Akhmanova, 2016; Vazquez-Pianzola and Suter, 2012). The founding member of this protein family, *Drosophila* BicD, was identified because of its essential role during oogenesis and embryo development where it transports mRNAs which control polarity and cell fate (Bullock and Ish-Horowicz, 2001; Suter and Steward, 1991; Suter et al., 1989; Wharton and Struhl, 1989). This process is mediated by its binding to the RNA-binding protein Egalitarian (Egl) (Dienstbier et al., 2009; Mach and Lehmann, 1997). In the meantime, BicD and its orthologs have been shown to control a diverse group of microtubule transport processes through binding to different cargoes or adaptor proteins (Hoogenraad and Akhmanova, 2016; Vazquez-Pianzola and Suter, 2012).

BicD can alternatively bind to Chc and this interaction facilitates Chc transport of recycling vesicles at the neuromuscular junctions and regulates endocytosis and the assembly of the pole plasm during oogenesis (Li et al., 2010; Vazquez-Pianzola et al., 2014). The best-known function of Chc is in receptor-mediated endocytosis as part of clathrin, a trimeric scaffold protein (called triskelion), composed of three Chc and three Clathrin light chains (Clc) (Brodsky, 2012). Aside from this, clathrin was shown to localize to mitotic spindles in mammalian and *Xenopus* cells (Fu et al., 2010; Royle et al., 2005) and to possess a non-canonical activity by stabilizing the spindle microtubules during mitosis (Royle, 2012). This function depends on clathrin trimerization and its interaction with Aurora A-phosphorylated Transforming Acidic Coiled-Coil protein 3 (TACC3) and the protein product of the colonic hepatic Tumor Overexpressed Gene (ch-TOG) (Booth et al., 2011; Fu et al., 2010; Lin et al., 2010; Royle and Lagnado, 2006; Royle et al., 2005). This heterotrimer forms inter microtubule bridges between Kinetochore-fibers (K-fibers), stabilizing these fibers and promoting chromosome congression (Booth et al., 2011; Royle et al., 2005). More recently, TACC3 and a mammalian homolog of Chc (CHC17) were shown to control the formation of a new liquid-like spindle domain (LISD) that promotes the assembly of acentrosomal mammalian oocyte spindles (So et al., 2019).

As *Drosophila* BicD forms complexes with both Chc and Dynein, two proteins with essential mitotic and meiotic activities, we set out to investigate possible *BicD* functions during cell division. BicD and Chc localize to the mitotic spindles and centrosomes, but also to the female tandem meiotic II spindles. Reduced BicD levels revealed that BicD is needed to correctly localize D-TACC, clathrin, and Msps (ch-TOG homolog) to the meiosis II spindles and for normal progression of meiosis II. BicD is required for the normal metaphase arrest of polar bodies after meiosis II completion and our results suggest that this activity is mediated by BicD’s role in localizing the spindle assembly checkpoint (SAC) components. Interestingly, *BicD* and *tacc* are also needed for pronuclear fusion after fertilization.

## RESULTS

### BicD and its cargo clathrin localize to centrosomes and spindles during mitosis and to tandem spindles in meiosis II

Completion of female meiosis and the first mitotic cycles depend on the correct spindle formation in the egg and the developing embryo. Maternally expressed genes provide all the proteins that control these processes, and their inactivation leads either to maternal effect lethality or female sterility. Indeed, *BicD* loss-of-function mutants are female sterile because they do not produce oocytes (Ran et al., 1994). This is an obstacle for studying the role of *BicD* in the maternally controlled early mitotic divisions of the embryo. Our laboratory has developed the *BicD*^*mom*^ females, a method to overcome *BicD* mutant female sterility (Swan and Suter, 1996). In *BicD*^*mom*^ females, BicD is provided from an inducible promoter that can be turned off once oocyte fate is established. Around 3-4 days after shutting down *BicD, BicD*^*mom*^ ovaries contain egg chambers devoid of BicD and few of them develop into eggs (Swan and Suter, 1996; Vazquez-Pianzola et al., 2014). Using this strategy, we observed that the eggs laid by *BicD*^*mom*^ females did not develop but arrested during stage 1 with phenotypes that required a more detailed analysis (see below) (Fig. S1A and SM1-4). This suggests that *BicD* is also essential downstream of oocyte differentiation to complete meiosis and progress through the early mitotic divisions.

We analyzed BicD localization during mitosis in methanol-fixed embryos. Methanol fixation dissolves the cytosolic pool of BicD, making insoluble pools of the protein more apparent. Surprisingly, during the syncytial divisions, BicD was detected on the centrosomes where it colocalized with the pericentrosomal marker centrosomin (Cnn), as well as on mitotic spindles (Fig. 1A,a,b, Fig. S1B). During cellularization, BicD was additionally enriched at the plasma membrane (Figure 1Ac). To confirm the specificity of the BicD antibody, we additionally analyzed embryos expressing BicD::GFP by immunofluorescence after staining them with anti-GFP. Both staining patterns were highly similar (Fig. 1B and Fig. S1C).

**Figure 1.**
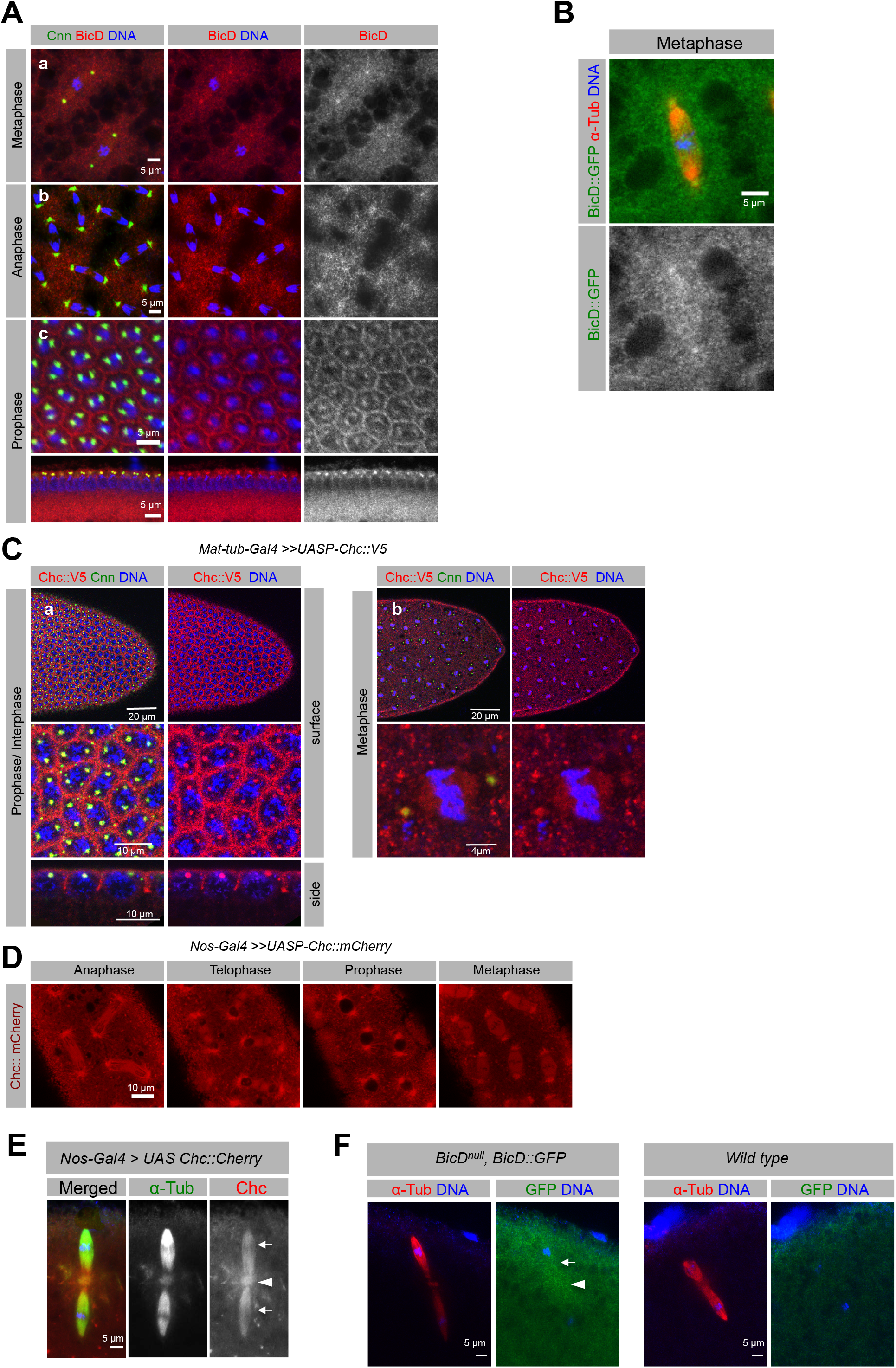
Dynamic localization of BicD and Chc with enrichment at embryonic spindles and centrosomes. **(A)** BicD (red), colocalized with the pericentrosomal marker Cnn (green) and it is also present at metaphase spindles. Wild-type embryos in nuclear cycle 10 (NC10) (a), NC12 (b) and during cellularization (NC14) (c) are shown. **(B)** *BicD*^*null*^, *BicD::GFP* stained with anti-GFP (green) and anti-α-tubulin (red) antibodies. An NC10 embryo is shown. **(C)** Chc::V5 expressing embryos stained for V5 (red) and Cnn (green). A cellular blastoderm (a) and an embryo in metaphase of NC13 are shown (b). **(D)** Live imaging of Chc::mCherry expressing embryos. **(E)** Chc::mCherry, detected with anti-mCherry antibodies (red), is present along both MII tandem spindles (arrows) and at the central aster (arrowhead). **(F)** During MII, BicD::GFP (detected with anti-GFP antibodies) is enriched in the region of the tandem spindles (arrow marks the most superficial spindle) and the central aster (arrowhead) above cytoplasmic levels. Embryos in E-F were also stained for anti-α-tubulin (red in E, green in F). In A-C, E-F, DNA was also stained with Hoechst (blue).

Chc is also present at the mitotic apparatus in vertebrates (Fu et al., 2010; Royle et al., 2005) and *Drosophila* Chc is transported by BicD (Li et al., 2010; Vazquez-Pianzola et al., 2014). Because available antibodies against Chc do not work well for immunostaining (Li et al., 2010), we analyzed the expression of the Chc in *Drosophila* embryos using tagged, yet functional, Chc fusions (Vazquez-Pianzola et al., 2014). Both, immunostaining of different tagged Chc proteins (Fig. 1C, Fig. S2A, Fig. S2B) and live imaging of embryos expressing fluorescent Chc (Fig.1D, Fig. S2C) revealed that Chc, like BicD, was enriched at the centrosomes and pericentrosomal regions during the entire cell cycle where it also co-localized with Cnn (Fig. 1C and Fig. S2B). During mitosis, Chc associates in addition with the mitotic spindles (Fig. 1C,D and Fig. S2C). This pattern of localization was not an overexpression artifact, because the localization of a Flag-tagged Chc expressed under its own promoter in a *Chc* null background, showed the same localization pattern (Fig. S2B). Moreover, similar to BicD, Chc was also enriched during cellularization near the plasma membrane between the nuclei, probably marking the sites where endocytic vesicles start to form (Fig. 1C) (Sokac and Wieschaus, 2008). A similar overall localization pattern was observed for the *Drosophila* Clathrin light chain (Clc) during embryogenesis (Fig. S2D). Interestingly, Chc was not only present at the mitotic spindles but also localized to the first spindles observed in the laid eggs, the female meiotic II tandem spindles. Chc signal was detected on both tandem spindles and the central aster (Fig. 1E). Assessing the localization of BicD in freshly laid eggs was challenging because of the high levels of BicD in the cytoplasm. Using the anti-BicD antibody, we observed a specific BicD signal, but this was at the same level in the meiotic spindle region as in the cytoplasm (Fig S1D). However, the presence of BicD at the meiotic spindle was more clearly detected with the anti-GFP antibody in *BicD::GFP, BicD*^*null*^ eggs. Similar to Chc, BicD::GFP was enriched above cytoplasmic levels on the meiotic tandem spindles and the central aster (Fig. 1F, and Fig. S1E,E’).

We conclude that BicD and Chc associate with the mitotic spindles and the centrosomal region during mitosis and with the tandem spindles and the central aster during female meiosis II. Thus, the localization of the Chc/Clc complex to the mitotic apparatus appears to be conserved between *Drosophila* and mammalian cells (Royle et al., 2005). Interestingly, mammalian BicD1 and BicD2 are present at the centrosomes in mammalian cells as well (Fumoto et al., 2006). Therefore, our data suggest that *BicD* might play a yet unidentified, but evolutionarily conserved role at the mitotic/meiotic spindles.

### deGradFP knockdown of BicD::GFP reveals a novel, essential role for BicD during early embryogenesis

Although *BicD*^*mom*^ flies lay embryos with mitotic defects, the number of eggs they lay is too small for phenotypic analyses. Thus, we designed a strategy to knock down the BicD protein directly in young embryos (stage 1) using the deGradFP technique (degrade Green Fluorescent Protein), a method to target fusion proteins with GFP for destruction or inactivation (Caussinus et al., 2011). For this, we took advantage of the functional, genomic *BicD::GFP* construct (Paré and Suter, 2000). Additionally, we constructed a deGradFP system specifically active during embryogenesis but not oogenesis by expressing the deGradFP (NSlmb-vhhGFP4) from a *hunchback* (*hb*) minimal maternal promoter coupled with the *bcd* 3’-UTR (Fig. 2A). The *hb* promoter is active during late oogenesis such that the *deGradFP* mRNA will be loaded into eggs and embryos. The *bcd* 3’ UTR promotes mRNA localization to the anterior pole of the oocyte and egg, and allows translation only upon egg activation in freshly laid eggs (Berleth et al., 1988; Driever and Nüsslein-Volhard, 1988; Sallés et al., 1994). We refer to this construct as *hb-deGradFP* (Fig. 2A). We corroborated the enrichment of the *deGradFP* mRNA in the anterior region of the embryo from egg-laying until before cellularization (Fig. 2B). Immunostaining experiments to detected *hb-deGradFP* expression revealed that the Vhh-GFP was distributed throughout the embryo and did not form an A-P gradient (Fig. 2C). It thus appears that the deGradFP is stable and can move to the rest of the embryo. Our new *hb-deGradFP* tool should therefore be useful to degrade GFP fusion proteins in entire young embryos.

**Figure 2.**
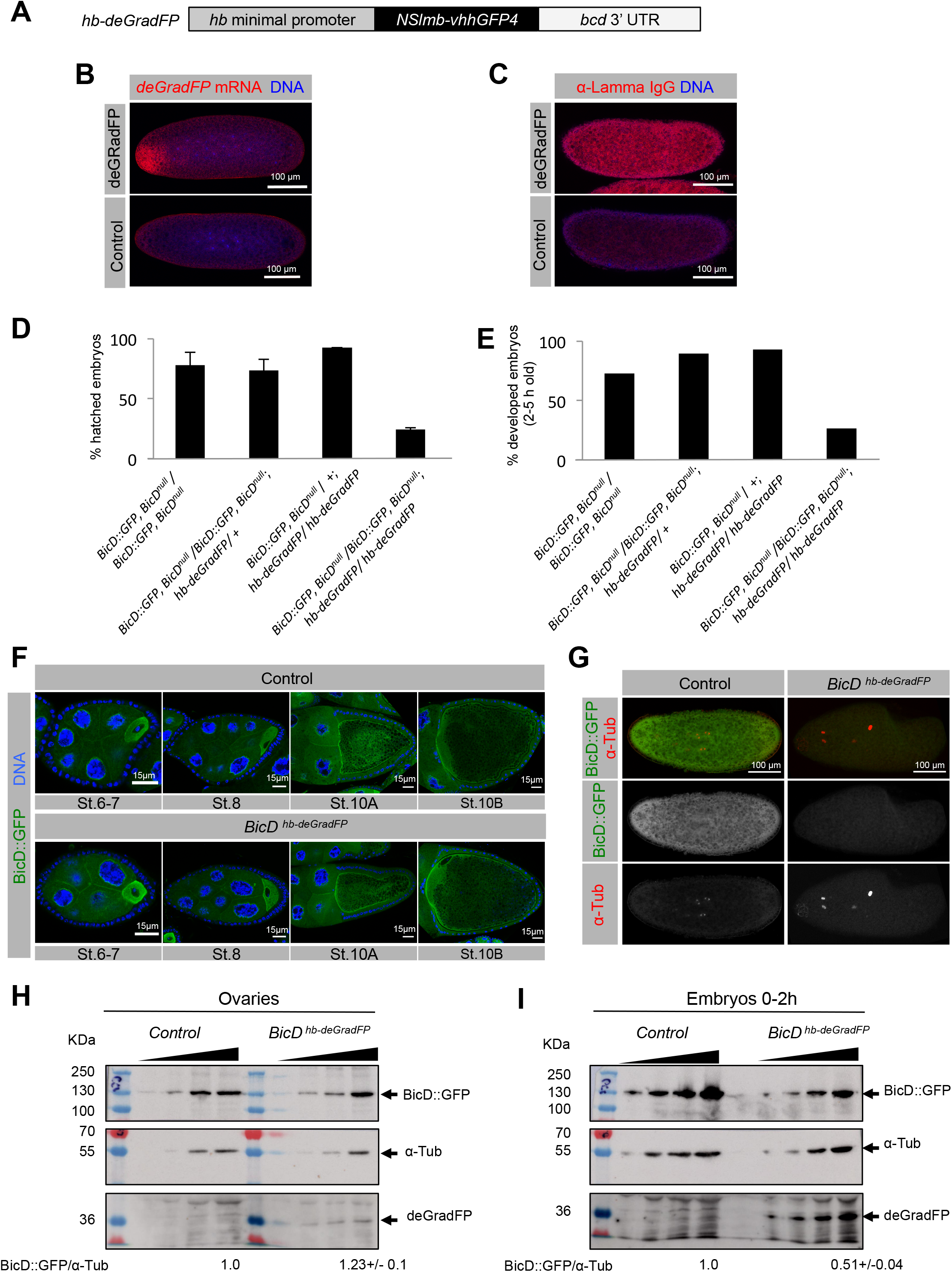
*hb-deGradFP* degrades BicD::GFP specifically in freshly laid eggs, causing early developmental arrest. **(A)** Scheme depicting the construct expressed in flies. **(B)** RNA FISH detects the *deGradFP* mRNA (red) enriched in the anterior region of freshly laid embryos. **(C)** The deGradFP construct (red signal) is ubiquitously present in young embryos. Wild-type embryos without the *deGradFP* transgene were used as negative controls in B and C. **(D)** Percentage of embryos of the depicted maternal genotype that hatched as larvae. **(E)** Percentage of normally developed embryos laid by the indicated mothers. **(F-I)** Ovaries and embryos produced from control (*BicD::GFP, BicD*^*null*^) and *BicD*^*hb-deGradFP*^ females were used. **(F)** Shows BicD::GFP fluorescence signal (green). **(G)** BicD::GFP was detected using anti-GFP antibodies (green). Embryos were staged by anti-α-tubulin staining (red). **(H-I)** Western blots showing BicD::GFP and deGradFP levels in ovaries **(H)** and 0-2h old embryos **(I)**. Increasing amounts of cytoplasmic extracts were loaded for each sample (loading control: α-tubulin). Relative abundance of BicD::GFP/α-tubulin, normalized to the control, was calculated for the last 2 lanes. Hoechst (blue) shows the DNA in IF experiments.

A previously described *BicD::GFP* construct rescued the sterility phenotype and the embryonic developmental arrest of *BicD* loss-of-function mutant females (*BicD::GFP, BicD*^*null*^ homozygous females) (Paré and Suter, 2000) (Fig. 2D-E). However, when these females expressed 2 copies of the *hb-deGradFP* construct (*BicD::GFP, BicD*^*null*^; *hb-deGradFP* homozygous females), 75% of their progeny failed to developed into late embryonic stages and did not hatch into larvae, showing, instead, arrest in meiosis or the first mitotic divisions as observed in embryos laid by *BicD*^*mom*^ females (Fig. 2D-E, Fig. SM5-7. In the following, we will refer to these progeny and their mothers as *BicD*^*hb-deGradFP*^. Embryos laid by females that expressed two copies of the *hb-deGradFP*, but also a wild-type *BicD*^*+*^ (*BicD::GFP, BicD*^*null*^/+(CyO); *hb-deGradFP*), did not show developmental problems, indicating that high levels of *hb-deGradFP* expression are not deleterious for development on their own (Fig. 2D-E). As assayed by immunofluorescence and Western blotting, *BicD*^*hb- deGradFP*^ females displayed normal ovaries and their BicD::GFP protein levels were not noticeably reduced in the ovary (Fig. 2F-H), confirming that the *hb-deGradFP* construct is not active during oogenesis. In contrast, in young *BicD*^*hb-deGradFP*^ embryos, the BicD::GFP signal was clearly reduced in the entire embryo (Fig. 2G) and BicD::GFP protein levels were downregulated by 50% (Fig. 2I). BicD::GFP is expressed at levels comparable to the wild-type BicD protein (Fig. S3A). However, 50% reduction in the levels of BicD::GFP protein detected by WB produces already visible phenotypes, while eggs from heterozygous females (*BicD*^*null*^/+) that have also a 50% reduction of BicD compared to wild-type embryos (Fig S3A) develop normally (97.3 ±1.03 % embryos hatched into larvae). Although alternative explanations for these differences are possible, we think that the deGradFP might bind BicD::GFP, functionally inactivating the protein before sending it to degradation.

As intended, the *hb-deGradFP* construct is expressed much higher in young embryos than in ovaries, although low levels of expression were also seen in ovaries (Fig. 2H, I) and even when these were dissected in the hypotonic Robbs medium that is normally used to avoid oocyte activation (Fig. S4A). Although we cannot rule out that some old egg chambers are activated during dissection due to physical activation, it is also possible that the *bcd* 3’-UTR sequence is not sufficient to fully control translation in the context of the maternal *hb* promoter and its 5’-UTR. Stage 14 (S14) oocytes are normally arrested in Metaphase I of meiosis I until the oocyte becomes activated by its passage through the oviduct. Because BicD is also present at the MI spindles in S14 oocytes (Fig. S4B), we further assessed the specificity of the early embryonic effect of *BicD*^*hb-deGradFP*^ by looking for evidence that *BicD*^*hb- deGradFP*^ causes meiosis I defects in oocytes (Fig. S4C-D). However, *BicD*^*hb-deGradFP*^ S14 oocytes displayed no evident problems in spindle formation or chromosome alignment in meiotic metaphase I, suggesting that the early embryonic arrest observed in *BicD*^*hb-deGradFP*^ individuals is not due to earlier meiotic defects during late oogenesis.

### *BicD* is required for the cell cycle arrest of the male and the female meiotic products, and for pronuclear fusion

To learn more about the novel BicD function in the earliest phase of embryonic development (Fig. 3) and to pinpoint the developmental stage, we collected fully viable control embryos and *BicD*^*hb-deGradFP*^ embryos over a 30 min period, let them develop for another 30 min (30-60 min old collections) and analyzed them for developmental defects (Fig. 3A). Twenty-five minutes after the eggs were laid, control embryos finished the 2^nd^ mitotic division and contained at least four zygotic nuclei. At this early stage, normal zygotic nuclei reside in the interior of the embryo and the three remaining female meiotic polar body products (mostly fused into one or two rosette-shaped nuclei) reside at the embryonic surface. Indeed, most embryos laid by control mothers (*BicD::GFP, BicD*^*null*^ or *BicD::GFP, BicD*^*null*^ / + (CyO); *hb-deGradFP)* developed normally, displaying more than four zygotic nuclei with normally looking mitotic spindles (Fig. 3A-B). However, embryos laid by *BicD*^*hb-deGradFP*^ mothers were arrested mostly at earlier stages, displaying spindle-like structures that appeared abnormal (Fig. 3A, C). Even though they contained mainly centrally located dividing nuclei, around 25% of these embryos were classified as “arrested with centrosomes” because they were positive for Cnn staining (Fig. 3A, an example in Fig 3Ca). In this category, we found embryos that contained at least one spindle displaying clear and sometimes fragmented staining for the centrosomal marker centrosomin (Cnn) at the spindle poles. Additionally, they frequently also displayed “free centrosomes” marked by Cnn signals associated with α-tubulin, but without a complete spindle and without DNA (example in Fig. 3Ca1). Embryos displaying acentrosomal spindles but containing free centrosomes were also scored into this category. Another 35% of the *BicD*^*hb-deGradFP*^ embryos possessed one or more internal acentrosomal spindle and all these spindles were negative for Cnn staining and were classified as “arrested, acentrosomal” (Fig. 3A, Cb).

**Figure 3.**
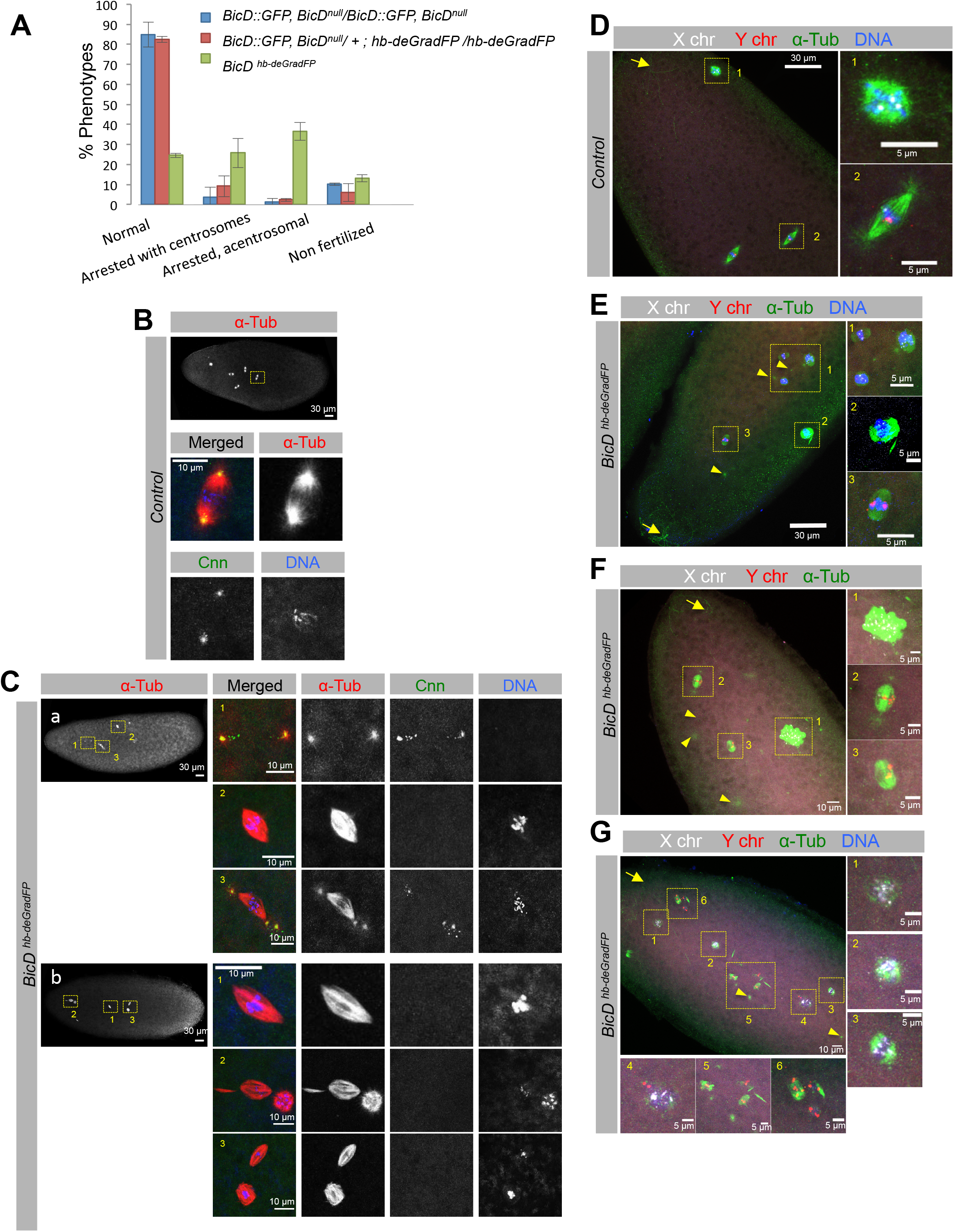
Fertilized *BicD*^*hb-deGradFP*^ embryos arrest at the start of embryogenesis with abnormal spindles, over-replicated female meiotic products, and without pronuclear fusion. **(A-C)** 30-60 min old embryos with the indicated maternal genotypes stained for α-tubulin (red), the centrosome marker Cnn (green), and DNA (blue). **A)** Percentage of embryos displaying the different phenotypes. The genotypes of their mothers are indicated. Embryos were characterized as unfertilized when no internal nuclei were seen and the 4 polar bodies, arrange in rosette-like structures (mostly fused into a single one), were observed at the surface of the embryo. Two independent collections (n1 and n2) were analyzed. Error bars show the SD. For *BicD::GFP, BicD*^*null*^*/BicD::GFP, BicD*^*null*^ *n1=84, n2=82*. For *BicD::GFP, BicD*^*null*^*/+; hb-deGradFP/hb-deGradFP n1=55, n2=70*. And for *BicD*^*hb-deGradFP*^ *n1=42, n2=63*. (**B-C**) Examples of the observed phenotypes. **(B)** Normal nuclear divisions in a control embryo (from a *BicD::GFP, BicD*^*null*^ mother). **(C)** Overview of the spindles seen in *BicD*^*hb-deGradFP*^ embryos classified as “arrested with centrosomes” (a) or as *“*arrested, acentrosomal” (b). **(D-G)** DNA FISH with X (white) and Y (red) chromosomal probes and stained for α-tubulin (green) and DNA (blue). Arrows mark the sperm tails and arrowheads free centrosomes. **(D)** Example of a control wild-type male embryo in the 2nd mitotic metaphase. **(E-G)** Examples of arrested *BicD*^*hb-deGradFP*^ male embryos. Images are z-stack projections through the nuclei and yellow boxes mark the corresponding magnified nuclei.

Consistent with the fact that centrosomes are inherited from the father, embryos arrested and classified as “arrested with centrosomes” were mostly marked by the presence of the sperm tail (Fig. S5A-B). In contrast, embryos displaying “acentrosomal”-like spindles and no free centrosomes rarely displayed any of these sperm tail markers, indicating that they more likely represent unfertilized eggs with aberrant meiotic products (Fig. S5A-B). We then analyzed in more detail the phenotype of arrested fertilized *BicD*^*hb-deGradFP*^ eggs by detecting the presence of the X and Y chromosomes by DNA fluorescent *in situ* hybridization (FISH). Male embryos (marked by the presence of the Y) develop only from fertilized eggs. Male embryos from control mothers showed one dot-like signal for the X chromosome and one signal for the Y in each zygotic nucleus, and these nuclei were located in the interior of the embryo (Fig. 3D1). The three polar bodies, formed after the two meiotic divisions, normally fused into a single polar body that was marked by the presence of the 3 X chromosomes (Fig. 3D2). In contrast, all arrested male embryos laid by *BicD*^*hb-deGradFP*^ females had at least one internal spindle marked only by the presence of the Y chromosome and no X chromosome signal, indicating that pronuclear fusion failed to take place and that a spindle still formed from the paternal pronucleus (Fig. 3E-G). In most of these embryos, the male pronucleus underwent only one additional round of replication since two dots of the Y chromosome signal can be observed in the internal metaphase spindle (70 %, Fig. 3E3). Embryos with more than one paternal spindle and many Y chromosome signals were also observed at a lower frequency (30%, Fig 3F2-3,G4-6). Independent of the rounds of replication observed in the parental pronucleus, these arrested *BicD*^*hb-deGradFP*^ embryos also contained one or several acentrosomal nuclei marked frequently by the presence of several dots of X chromosomal signal, suggesting that the polar bodies did not arrest in metaphase II as they normally do but underwent several cycles of DNA replication instead (example in Fig. 3E1-2, 3F1, 3G1-4). These results show that *BicD* is required for the female and male pronucleus cell cycle arrest and for pronuclear fusion.

### BicD is needed for replication arrest of the polar bodies and for their rosette formation

To test for an essential function of *BicD* during the final phase of the meiotic divisions, we crossed *BicD*^*hb-deGradFP*^ and control females to sterile XO males, causing them to lay unfertilized eggs (Fig. 4). In wild-type unfertilized eggs, egg activation is triggered by passage through the oviduct, and this causes the eggs to complete meiosis II. In collections of 0-1h old unfertilized control eggs, we observed from 1 to 4 rosette-like nuclei. These represent intermediate stages of the fusion process of the four meiotic products, which ultimately fused to form a single, rosette-shaped nucleus in wild-type eggs indicating that eggs completed meiosis II. These rosette-shaped nuclei were also marked by the presence of a total number of 4 dots of X-chromosomal signal per egg, which arise from each of the four meiotic products (Fig. 4A, examples in Fig. 4B). In contrast, *BicD*^*hb-deGradFP*^ unfertilized eggs contained one to several nuclei forming spindle-like structures, mostly with the appearance of multipolar spindles (Fig. 4A). Additionally, their chromosomes did not create a rosette structure that would be typical for a metaphase arrested state. Instead, these nuclei displayed partially decondensed chromatin, an irregular shape, and they lacked the α-tubulin staining ring that surrounds the DNA rosette in control eggs (Fig. 4A,C). Furthermore, the X-chromosomal probe produced more than the normal four signal dots per egg, in eggs containing only one nucleus (Fig. 4C-c) and in eggs containing several nuclei and spindles (Fig. 4C-c’). We also observed eggs with more than four meiotic nuclei (like in Fig. 4Cc’), indicating that *BicD*^*hb-deGradFP*^ eggs show over-replication of the meiotic products. The probe used for the FISH experiments recognizes a repetitive region on the X chromosome present along 3 to 3.5 Mb. The fact that this probe detected more than four signals in each *BicD*^*hb-deGradFP*^ egg, and that these signals showed different brightness and sizes might also suggest that the DNA has become fragmented and/or more decondensed and that it did not arrest in a metaphase-like state as in the normal rosette structures. This replication, decondensation and/or fragmentation of the meiotic DNA was not restricted to the sex chromosomes because we observed analogously additional signals when using a probe for the 2^nd^ chromosome (Fig. S6). Altogether, these results indicate that BicD is required for both the replication arrest and the formation of the typical rosette-like structures of the polar bodies.

**Figure 4.**
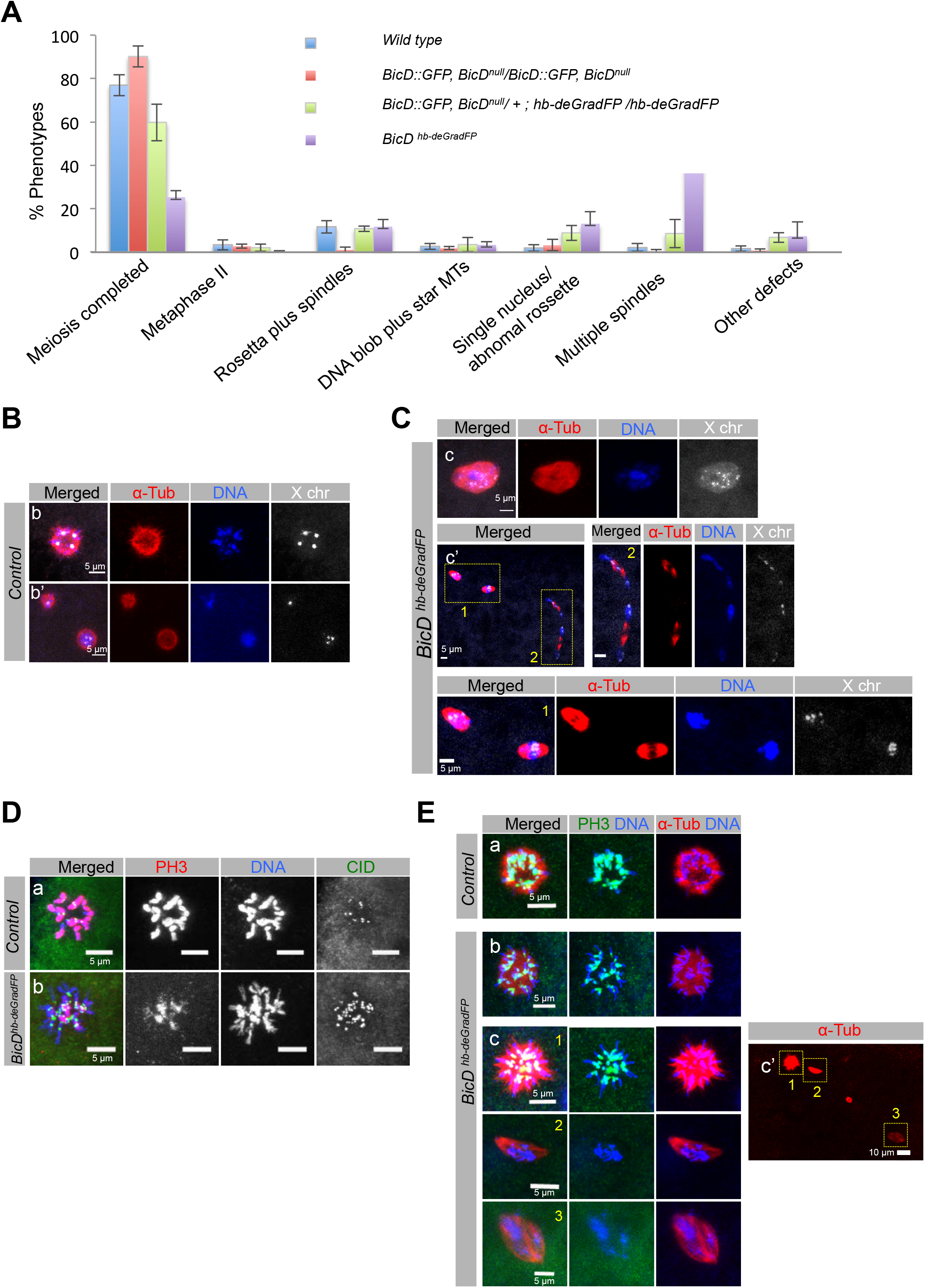
Female meiotic products fail to arrest in metaphase and undergo extra rounds of replication in unfertilized *BicD*^*hb-deGradFP*^ eggs. **(A)** 0-1 h old unfertilized eggs from the depicted mothers stained for α-tubulin (red) and DNA (blue) and classified as follows. 1) eggs that apparently completed meiosis normally (up to four rosette like structures that normally fuse into one single rosette); 2) eggs displaying a normal metaphase II tandem spindle; 3) eggs presenting 1-2 rosette like structures plus 1-2 spindles (probably representing normal meiosis intermediate products); 4) eggs with a big DNA blob surrounded by a star of microtubules (MT); 5) eggs containing one single nucleus/ big spindle; 6) eggs containing several spindles; 7) eggs with other defects. Most control eggs (from wild type, *BicD::GFP, BicD*^*null*^ or *BicD::GFP, BicD*^*null*^*/+; hb-deGradFP* mothers) completed meiosis normally. In contrast, many eggs from *BicD*^*hb-deGradFP*^ mothers contained multiple spindles and, to a lesser extent, one big spindle. Error bars show the SD of three to four independent egg collections (n1 to n4). For *wild type*, n1= 145, n2=95, n3=67; for *BicD::GFP, BicD*^*null*^*/BicD::GFP*, n1=92, n2=72, n3=147; for *BicD*^*null*^, *BicD::GFP/ +; hb-deGradFP /hb-deGradFP*, n1=68, n2=46, n3= 69, n4=67; for *BicD*^*hb-deGradFP*^, *n1=29, n2=45, n3=32* were scored. **(B-C)** Eggs from the same parents as in A) subjected to FISH to detect the X chromosome (white) and stained for α-tubulin (red) and DNA (blue). **(B)** Examples of control eggs (*BicD::GFP, BicD*^*null*^), where the meiotic products have fused into a single (b) or two rosette-like structures (b’), showing a total of four X chromosomes signals. **(C)** (c) *BicD*^*hb-deGradFP*^ egg with a single nucleus containing an abnormal rosette-like structure, partially decondensed chromosomes, and many dotted X chromosomal signals. (c’) *BicD*^*hb-deGradFP*^ egg with more than four meiotic products. Every nucleus contains many dotted X chromosomal signals (C1-2). **(D-E)** Unfertilized wild-type and and *BicD*^*hb-deGradFP*^ eggs stained for CID (green, marking centromeres), PH3 (red) and DNA (blue). 100% (12/12 confocal images) of the single fused rosette nuclei in control eggs showed strong PH3 staining along the entire chromosomes. Although these structures in *BicD*^*hb-deGradFP*^ eggs were also positive for PH3, in 90,9 % (10/11) of them the signal was only enriched at the pericentromeric region. 86% (6/7 confocal images) of the single fused rosette-like nuclei in control eggs showed normal CID staining. In contrast, 63.7 % (7/11) of these structures in *BicD*^*hb-deGradFP*^ eggs showed an increased number of CID-positive dots. **(E)** Eggs were stained for α-tubulin (red), PH3 (green), and DNA (blue). Images are z-stack projections through the nuclei and yellow boxes mark the corresponding magnified nuclei.

### Role of BicD in SAC and metaphase arrest of female meiotic products

After meiosis II is completed, *Drosophila* polar bodies remain arrested in a metaphase-like state. In wild-type unfertilized eggs, the four meiotic products, which fused into a single rosette, showed strong signal for the mitotic marker Phospho-Histone 3 (PH3) along the entire chromosomes, indicating that they were arrested in a metaphase-like state (Fig. 4Da-Ea). In contrast, in *BicD*^*hb-deGradFP*^ eggs showing one rosette-like structure, indicative of meiosis completion, the PH3 staining was not localized along the entire chromosomes but only enriched at the pericentromeric region (Fig. 4 Db-Eb). Moreover, their rosette-like structures showed an increased number of CID-positive dots suggesting that female meiotic products underwent extra rounds of replication/endoreplication (compare Fig. 4Da with 4Db).

Rosette-like structures in *BicD*^*hb-deGradFP*^ eggs did also not form the typical tubulin ring surrounding the chromosomes observed in wild-type eggs and, interestingly, DNA extended beyond this tubulin ring and this DNA was negative for PH3 staining (Fig. 4Ea, b, c). In the normal situation, Histone H3 phosphorylation starts in pericentromeric heterochromatin regions at the onset of mitosis and then spreads along the entire length of chromosomal arms, reaching its maximal abundance during metaphase. This is then followed by a rapid decrease upon transition to anaphase (Sawicka and Seiser, 2012). Thus, PH3 staining confined to the pericentromeric region in *BicD*^*hb-deGradFP*^ polar bodies suggests that these nuclei are not properly arrested or are released from metaphase arrest. Furthermore, *BicD*^*hb-deGradFP*^ eggs possessing several meiotic products (Fig. 4Ec,c’) showed that not all of these nuclei were positive for PH3 staining (Fig. 4Ec2-3,c’), further strengthening the idea that these nuclei are over-replicating due to a failure to arrest in metaphase.

The metaphase arrest of polar bodies depends on the activation of the spindle assembly checkpoint (SAC) pathway (Défachelles et al., 2015; Fischer et al., 2004; Pérez-Mongiovi et al., 2005). We, therefore, analyzed the localization of two well-conserved orthologs of the SAC pathway, BubR1, and Mad2 (Fig. 5). These proteins associate with unattached kinetochores and, in the case of BubR1, also to kinetochores lacking tension. By inhibiting the anaphase-promoting complex (APC/C) they are essential to maintain the metaphase arrest. BubR1 was clearly present at polar body kinetochores in the wild type (100%, n=23) and in the control BicD::GFP rescued eggs (93%, n=27) (Fig. 5Aa). However, in 43% of the *BicD*^*hb-deGradFP*^ eggs, the meiotic products failed to recruit BubR1 to the kinetochores (n=35). The absence of BubR1 from the polar body kinetochores was observed in eggs where polar bodies were fused into a single rosette (Fig. 5Ab) and in eggs showing many additional meiotic products (Fig. 5Ad, d’). The rest of the *BicD*^*hb-deGradFP*^ eggs showed either normal BubR1 recruitment (40%) or only a weak signal for kinetochore BubR1 (17%, Fig. 5Ac). These data indicate that the failure to activate or maintain the metaphase arrest of polar bodies in the absence of BicD is probably due to a failure to recruit the SAC components to the kinetochores or maintain their association. The fact that half of the polar bodies still recruited SAC components might also suggest that these nuclei are cycling in and out of the metaphase arrest, duplicating their chromosomes. Similar results were obtained for Mad2, where 69% of the *BicD*^*hb-deGradFP*^ eggs analyzed did not show recruitment of Mad2 to the polar bodies (n=48; Fig. 5B). Altogether, these results suggest that in *BicD*^*hb-deGradFP*^ eggs, the meiotic II products do not correctly arrest in metaphase due to a failure to activate or maintain the SAC.

**Figure 5.**
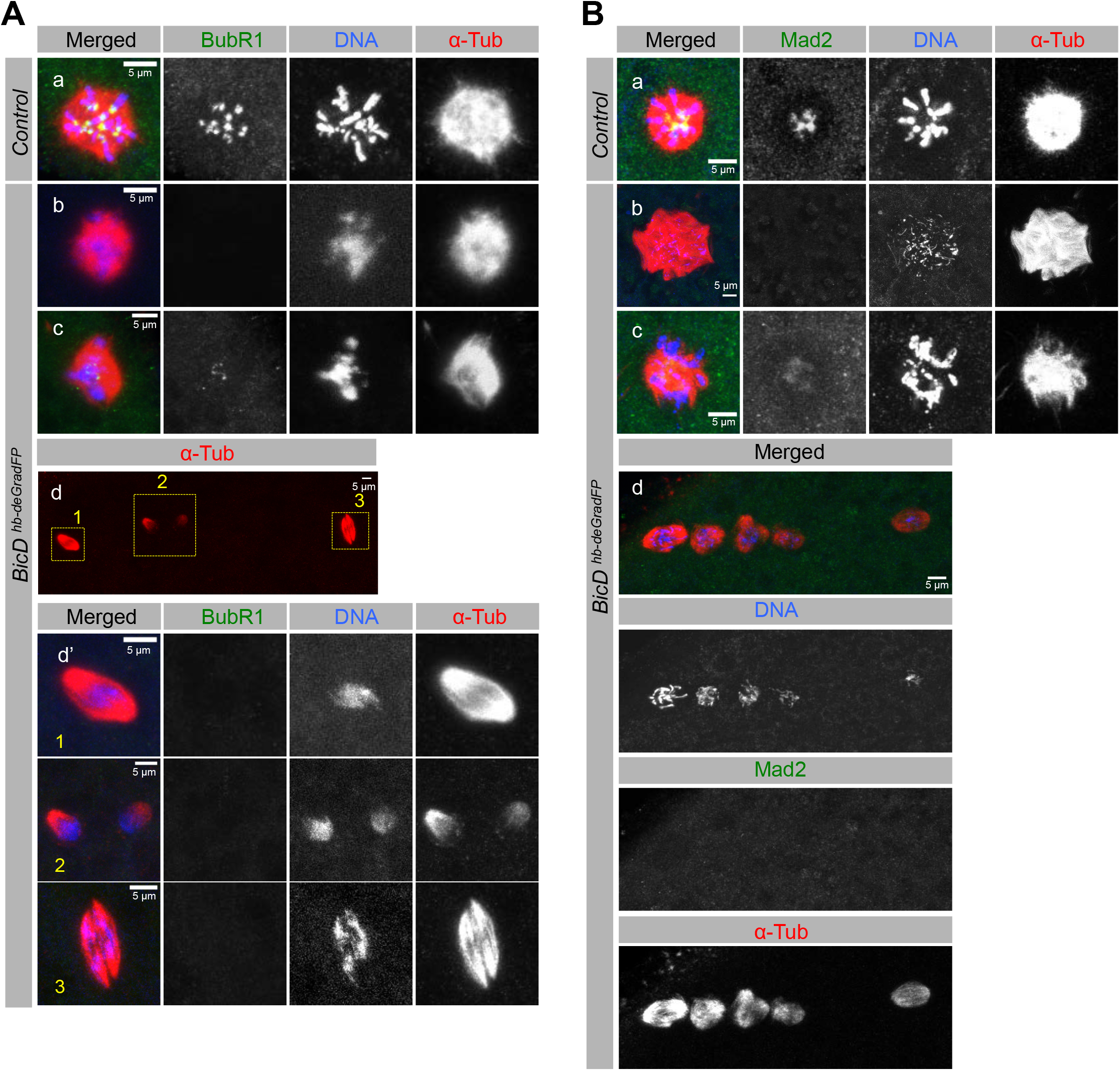
BicD contributes to the recruitment of the SAC components BubR1 and Mad2 to the female meiotic products. 0-1 h old unfertilized eggs stained for α-tubulin (red), DNA (blue) and BubR1 **(A)** or Mad2 **(B)** (green). (**A) a)** Wild-type meiotic products fused into a single rosette showed BubR1 staining at the kinetochores. **b)** *BicD*^*hb-deGradFP*^ egg containing a unique rosette-like polar body without BubR1 signals. **c)** *BicD*^*hb-deGradFP*^ egg with only one polar body showing weak BubR1 signals. **d)** *BicD*^*hb-deGradFP*^ egg with several meiotic products that are negative for BubR1. **d’)** magnified images of the indicated nuclei in d). **(B) a)** Single polar body rosette structure in a control egg (*BicD::GFP, BicD*^*null*^) with Mad2 at the kinetochores of the polar body. **b)** *BicD*^*hb-deGradFP*^ egg where the unique rosette-like polar body lacks Mad2 signal. **c)** *BicD*^*hb-deGradFP*^ egg with only one polar body that stains only weakly for Mad2. **d)** In this *BicD*^*hb-deGradFP*^ egg, the female meiotic products have replicated and don’t stain for Mad2. Images are z-stack projections through the nuclei.

### Proper localization of D-TACC, Msps, and Clc at tandem meiotic spindles requires BicD

To pinpoint the first meiotic defects, we focused on the very early stages of meiosis II in eggs that were just released from the MI arrest. The analysis of Chc distribution proved difficult due to the lack of useful antibodies for immunostaining. Localization of tagged Chc was also irreproducible due to the high cytoplasmic signal and the fact that overexpressing tagged Chc in a background of reduced BicD levels increased the *BicD*^*hb-deGradFP*^ phenotype (see Fig. 7C-D). However, because the Chc partner Clc is also present at the mitotic spindles together with Chc (Fig.S2D), we followed Clc localization during meiosis II. Clc localized to the female MII tandem spindles and the central aster (Fig. 6A). Additionally, we detected an unusual, strong accumulation of Clc at the central aster relative to the levels observed along the MII spindles in *BicD*^*hb-deGradFP*^ compared to control spindles (Fig. 6A, A’). We then followed the localization of D-TACC and Msps (Mini spindles) because the spindle localization of their mammalian homologs is interdependent with Chc (Royle, 2012; Royle et al., 2005). In wild-type MII and anaphase II (AII) spindles, D-TACC and Msps were present on tandem spindles, and they were enriched at the central aster (Fig. 6B, C arrowheads). In both metaphase II and anaphase II, D-TACC is additionally weakly enriched at the spindles’equator, where the MT plus ends are located (Fig. 6B, arrows). On the other hand, Msps was clearly enriched along both arms of the tandem spindles but stained more strongly the minus ends at the spindle poles (Fig. 6C, arrows). In *BicD*^*hb-deGradFP*^ embryos, D-TACC and Msps localization along the tandem spindles was generally clearly reduced compared to the control spindles (Fig. 6B, C). However, like in control spindles, D-TACC and Msps still showed enrichment at the central aster in *BicD*^*hb-deGradFP*^ embryos (Fig. 6B, C). Accordingly, the signal intensity for D-TACC and Msps along the most superficial tandem spindle relative to the signal intensity observed in the central aster was significantly reduced in *BicD*^*hb-deGradFP*^ MII spindles (Fig 6B’, C’). Thus, these results suggest that BicD is needed to properly localize clathrin, D-TACC and Msps along the meiotic II spindles.

**Figure 6.**
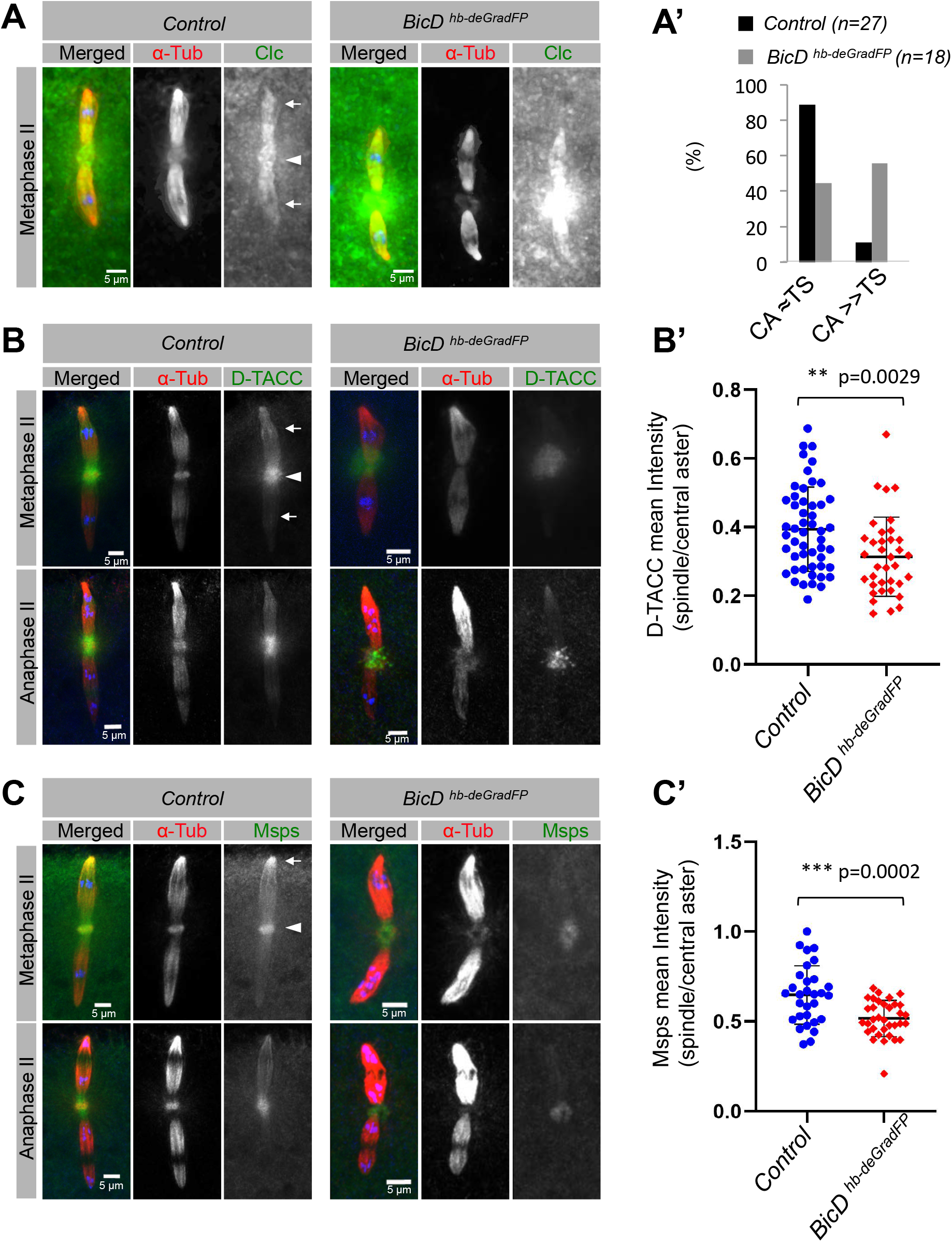
Proper localization of D-TACC, Msps, and clathrin at the MII tandem spindles requires BicD. 5 min egg collections of controls (laid by homozygous *BicD::GFP, BicD*^*null*^ (B-C) or wild-type mothers for (A) and *BicD*^*hb-deGradFP*^ embryos stained for α-tubulin (red) and either Clc **(A)**, D-TACC **(B)** or Msps **(C)** (green). DNA is shown in blue. **(A)** Maximum intensity projection images subjected to depth correction (see Methods) are shown. Clc hyper-accumulates at the central aster (arrowhead) compared to the signal at the spindles (arrows) *in BicD*^*hb-deGradFP*^eggs. (A’) Percentage of embryos showing similar signal intensity along the tandem spindles (TS) and the central aster (CA) (CA ≈TS), and embryos with hyper-accumulation of Clc at the central aster (CA>>TS) is shown for both genotypes. n= sum of embryos scored for each genotype in two independent experiments. Two independent researchers performed the qualitative scoring blindly. **(B-C)** Maximum intensity projection images are shown. The spindle shown on top is the more superficial one in the embryo and its signal is stronger because the signal from the spindle below is often masked by the cytoplasmic signal above. D-TACC and Msps localized to the tandem spindles and to the central aster (arrowheads). Msps is also enriched at the minus end poles of the tandem spindles in control embryos (arrow in C). Compared to the signal intensity in the central aster, D-TACC and Msps signals on the *BicD*^*hb-deGradFP*^ spindles are reduced. **(B’, C’)** The mean fluorescent signal intensity ratio (signal in the most superficial spindle vs signal in central aster) for D-TACC (B’) and Msps (C’) was quantified for each imaged MII and AII spindle. Data are presented as mean ± SD. Statistical significance and p values were determined by the Student’s unpaired test. (B’) Control: n=51, *BicD*^*hb-deGradFP*^: n=36; 3 experiments. (C’) Control: n=30, *BicD*^*hb-deGradFP*^: n=34; 2 experiments.

**Figure 7.**
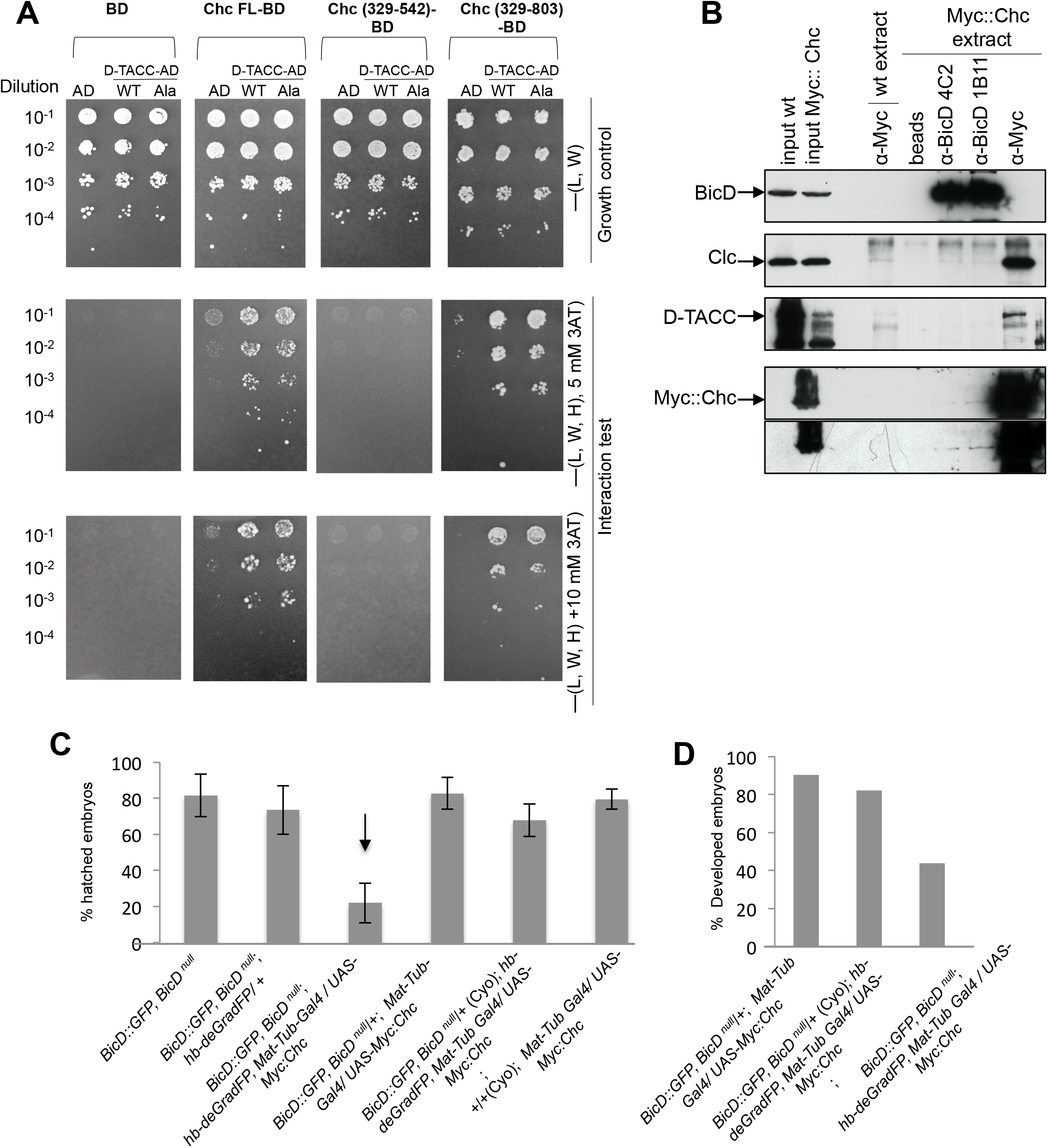
*Drosophila* Chc binds to D-TACC through the same domain it interacts with BicD, and *BicD* and *Chc* genetically interact in embryos. **(A)** Yeast two-hybrid interaction tests. Full-length (FL) and partial Chc fragments (Chc^329-542^ and Chc^329-803^), respectively, were fused to the DNA binding domain (BD). Full-length D-TACC wild type (WT) and *D-TACC*^*S863A*^ were fused to the activator domain (AD). Empty vectors served as negative controls. **(B)** IP of total embryo extracts expressing a Myc-tagged Chc fusion protein (Myc::Chc). Antibodies used for the IPs are indicated on top. Beads alone and anti-Myc pull-downs in wild-type (wt) embryo extracts were done as negative controls. Western blots of the precipitated material were tested for the presence of BicD, Clc, D-TACC and Myc::Chc. 0.15% of the cytoplasmic extract used for each IP was loaded as input. Bottom panel shows a longer exposure of the Myc::Chc blot. **(C-D)** Maternal genotypes are indicated. Percentage of embryos that hatched into larvae **(C)** and of developing embryos **(D)**.

### Evidence for mutually exclusive complexes of BicD/Chc and D-TACC/Chc

In mammals, Chc interacts directly with TACC3, and the interaction depends on the phosphorylation of Ser^552^ of TACC3 by Aurora A (Fu et al., 2010; Lin et al., 2010). Since D-TACC and clathrin fail to properly localize on *BicD*^*hb-deGradFP*^ MII spindles, we further investigated whether the *Drosophila* homologs of these proteins can interact with each other and with BicD using a yeast 2-hybrid (Y2H) assay and co-immunoprecipitation experiments (Fig. 7A-B). Indeed, full-length *Drosophila* Chc interacts with D-TACC in both assays.

The minimal region of mammalian isoform Chc22 that interacts with TACC3 consists of amino acids 331-542 (Lin et al., 2010). Whereas the corresponding region of *Drosophila* Chc^329-542^ did not interacted with D-TACC in our assays (Fig. 7A), this region with a C-term extension (Chc^329-803^) interacted with D-TACC almost as strongly as the full-length protein (Fig. 7A). Interestingly, Chc^329-542^ was also unable to bind BicD (Fig. S7A), but Chc^329-803^ had been shown previously to contain the minimal BicD binding region (Li et al., 2010) (Fig. S7B). Chc^329-803^ interacted in our assays with the BicD CTD (C-terminal domain), a truncated version of BicD that normally interacts stronger with its cargoes because it does not contain the BicD fold-back domain (Fig. S7B) (Li et al., 2010). Since the sequences required for the binding of *Drosophila* Chc to D-TACC and to BicD are overlapping, Chc/BicD and Chc/D-TACC might only exist as mutually exclusive protein complexes. However, because Chc normally acts as a trimer, higher-order complexes of the three Chc molecules could theoretically contain BicD or D-TACC, or both. Surprisingly, the interaction of Chc with D-TACC was not abolished when the conserved Ser residue targeted by the Aurora A kinase was mutated to Alanine (Ser^863^ in D-TACC, corresponds to Ser^552^ of the human protein, Fig. 7A, Fig. S7C). Nevertheless, the phosphomimetic substitution Ser^863^>Asp in D-TACC increased its interaction with full-length Chc (Fig. S7C). These results suggest that phosphorylation of Ser^863^ may not be a prerequisite for the interaction of full-length D-TACC with Chc at least in the Y2H system. Still, it could enhance the interaction during mitosis and meiosis when the kinase is active.

The interaction of D-TACC with full-length Chc was observed under very stringent Y2H conditions (medium -L, -W, -H, -a; Fig. S7C). In contrast, the interaction between Chc and BicD was only visible under less stringent conditions (medium -L, -W, -H +3 mM 3AT; Fig. S7A, (Cagney et al., 2000)), suggesting that Chc interacts stronger with D-TACC than with BicD in this system. This is also supported by the immunoprecipitation experiments using embryo extracts. Using extracts expressing a Myc::Chc fusion protein, Myc::Chc was observed at low levels in BicD IPs. In contrast, the presence of BicD was not detected in the reverse IPs (with anti-Myc antibodies) as previously reported (Fig. 7B) (Vazquez-Pianzola et al., 2014). This is conceivable due to their weak interaction or because the Myc tag interferes with the binding sites. In contrast, D-TACC strongly co-immunoprecipitated with Myc::Chc. On the other hand, no interaction was observed between D-TACC and BicD either by IP or using the Y2H system (Fig. 7B, S7D).

### High levels of *Chc* enhance the embryonic arrest phenotype of *BicD*^*hb-deGradFP*^ embryos

*BicD::GFP, BicD*^*null*^ females, expressing only one copy of the *hb-deGradFP* construct *(BicD::GFP, BicD*^*null*^; *hb-deGradFP-bcd 3’UTR/+)* laid embryos that show a reduction of BicD::GFP levels only by 30% on WB experiments, (Fig. S3B-C) and mostly developed normally and hatched into larvae (Fig. 2D-E, 7C-D). However, when these females also expressed a copy of a *Chc* transgene under the control of the strong maternal tubulin promoter, 80% of their progeny failed to hatch as larvae and, instead, arrested with the same abnormal nuclei phenotype observed in *BicD*^*hb-deGRadFP*^ embryos (Fig. 7C-D). Overexpression of this Chc construct alone did not produce visible phenotypes in embryos, suggesting that high levels of Chc in a BicD-reduced background are responsible for the phenotype.

### Like BicD, *d-tacc* is required for pronuclear fusion and cell cycle arrest of the male pronucleus and the female meiotic products

*d-tacc*^*1*^ mutant mothers lay eggs with reduced D-TACC protein levels and most of these embryos fail to develop (Gergely et al., 2000). Of the male progeny laid by *d-tacc*^*1*^*/ Df(3R)110* females, only 25.8% showed signals for the X and Y chromosomes in each of their zygotic nuclei, indicating that they had performed pronuclear fusion (Fig. S8A). The majority of them (74.2%) had at least one internal nucleus strongly marked for the Y chromosome and no X-chromosomal signal (Fig. S8 B-D,), even though, X chromosome signals strongly marked the remaining nuclei which represent the female meiotic products. These data show that in *d-tacc*^*1*^ embryos (Fig. S8), pronuclear fusion is strongly compromised. From these embryos that did not perform pronuclear fusion, 30.4 % showed normal cell cycle arrest of the female and male meiotic products (Fig. S8B). However, in the majority of them, the paternal pronucleus underwent additional divisions with no over-replication of female meiotic products (47.85 %, Fig. S8C). Only 21.7% *d-tacc*^*1*^ showed replication of the female meiotic and male derived nuclei (Fig. S8D). The lower penetrance of this female meiosis phenotype compared to the one observed in *BicD*^*hb-deGRadFP*^ eggs (70% of the arrested eggs), could be because *d-tacc*^*1*^ embryos produce high enough levels of D-TACC, sufficient to rescue the female meiosis phenotype (Gergely et al., 2000). BicD and D-TACC are therefore needed for pronuclear fusion and metaphase arrest of the male pronucleus and the female meiotic products. Interestingly, we also observed abnormal meiosis with delayed polar body extrusion in eggs from *C. elegans* worms fed with bacteria expressing dsRNA against *tac-1* (Fig. S9C). Moreover, pointing to a conserved role for these proteins in pronuclear fusion, we found that eggs from worms fed with dsRNA against *bicd-1, chc-1* and *tac-1* did not undergo pronuclear fusion either (Fig. S9A-E).

## DISCUSSION

### A useful strategy to study the effect of lethal or female sterile mutations in early embryogenesis reveals that *BicD* is required for meiosis II and pronuclear fusion

In this study, we found that BicD localizes to the female meiotic spindle during meiosis II, where it is present at the tandem spindles and the central aster. After fertilization, BicD also localizes to the mitotic spindles and the centrosomes. Since *BicD*^*null*^ mutants rarely survive and are sterile, we set up a strategy based on the deGDradFP technique to assess its role during the early embryonic divisions. Using this strategy, we were able to generate embryos with reduced levels of BicD at the very beginning of embryogenesis (*BicD*^*hb-deGradFP*^ embryos). Consistent with BicD localization at the female meiotic II spindles, we discovered that *BicD*^*hb-deGradFP*^ embryos arrest development displaying aberrant meiotic products and no pronuclear fusion. Combined with the CRIPSP-Cas9 strategy first to produce functional GFP tagged versions of the proteins of interest, the construct designed in this paper could be useful to study the role of other female-sterile and lethal mutations during very early embryonic development (Nag et al., 2018).

### Connecting BicD to the SAC pathway

In unfertilized *BicD*^*hb-deGradFP*^ eggs, the female meiotic products were not arrested in metaphase as it normally happens. Instead, they underwent additional rounds of replication. *BicD*^*hb-deGradFP*^ female meiotic products failed to recruit or maintain the recruitment of the SAC pathway components BubR1 and Mad2, which are normally present at the kinetochores of the female meiotic polar bodies in wild-type eggs. Interestingly, in *Drosophila* mutants for *Rod, mps1*, and *BubR1*, well-conserved orthologs of the SAC pathway, the polar bodies cannot remain in a SAC-dependent metaphase-like state and decondense their chromatin, too (Défachelles et al., 2015; Fischer et al., 2004; Pérez-Mongiovi et al., 2005). Furthermore, in these mutants, the polar bodies have been shown to cycle in and out of M-phase, replicating their chromosomes similarly to what we observed in *BicD*^*hb-deGradFP*^ eggs. Thus, it appears that BicD normally functions to localize these SAC components to induce and/or maintain the metaphase arrest of the polar bodies. Several mechanisms could explain the failure to maintain SAC activation observed in *BicD*^*hb-deGradFP*^ embryos. BicD might be needed to recruit the SAC components to kinetochores directly. On the other hand, during mitosis, dynein motors have been shown to transport SAC components along the microtubules away from kinetochores as a mechanism to trigger checkpoint silencing and anaphase onset (Basto et al., 2004; Howell et al., 2001; Wojcik et al., 2001). Since BicD binds dynein, we expect that if BicD acts as a linker to transport SAC components, it might be required to move them away from the kinetochores. If this does not happen, the SAC remains persistently activated. Whereas persistent SAC activation leads to metaphase arrest and delayed meiosis (D-Meiosis), at least during mitosis, this delay is known to be rarely permanent. Most cells that cannot satisfy the SAC, ultimately escape D-mitosis and enter G1 as tetraploid cells by a mechanism that is poorly understood (Rieder and Maiato, 2004). Thus, it is possible that in embryos with reduced *BicD* function, the SAC pathway is constantly activated, delaying meiosis. However, at one point, the nuclei might escape the metaphase II arrest, cycling in and out of M-phase, thereby replicating their chromosomes and de-condensing their chromatin. The fact that female meiotic products in *BicD*^*hb-deGradFP*^ embryos show no or only pericentromeric PH3 staining and that they are replicating, supports the notion that these nuclei are on an in-out metaphase arrest phase. This is also supported by the finding that the meiotic products in around half of the *BicD*^*hb-deGradFP*^ embryos failed to stain for the SAC components BubR1 and Mad2.

### Connecting BicD to the D-TACC, Msps, and clathrin complex

Chc is transported by BicD in oocytes and neurons and it shows a similar pattern of localization as BicD (Li et al., 2010; Vazquez-Pianzola et al., 2014). During cellularization, both are localized in a dotted pattern near the invaginating plasma membrane, probably marking the sites where endocytic vesicles are formed. This is consistent with the described role of BicD in regulating endocytosis in oocytes and neurons. During mitosis, Chc, its partner Clc, and BicD are enriched at mitotic spindles and centrosomes. Furthermore, clathrin, BicD, and the clathrin-interacting partners D-TACC and Msps localize to the tandem spindles and the central aster of the female meiotic II apparatus. The interaction of *Drosophila* Chc with D-TACC is conserved and Chc interacts through the same protein domain directly with D-TACC and BicD (Fig 7, (Fu et al., 2010; Lin et al., 2010)). Additionally, embryos laid by mothers overexpressing Chc in a background where BicD is reduced down to levels that do not produce visible phenotypes on their own, also arrest during early development. Moreover, BicD is needed for localizing normal levels of D-TACC, Msps, and clathrin throughout the tandem spindles relative to their localization at the central aster (Fig. 6). Thus, it appears possible that limited BicD activity in *BicD*^*hb-deGradFP*^ embryos is insufficient to localize clathrin, TACC, and Msps efficiently along the MTs of the spindle. During mitosis, impairment of MT motors, such as dynein, as well as interference with MT dynamics, persistently activates the SAC (Rieder and Maiato, 2004). Treatments that prevent the TACC/ clathrin complex from binding to the mitotic spindles affect K-fiber stability and persistently activate the SAC, too (Rieder and Maiato, 2004; Royle et al., 2005). As discussed before, this SAC hyperactivation would also explain the over-replication of female meiotic products observed in eggs with reduced BicD or D-TACC levels.

### Role of BicD in pronuclear fusion

BicD is also needed for zygotes to perform pronuclear fusion (Fig 3). Furthermore, its role in pronuclear fusion is conserved during evolution since *C. elegans* eggs depleted for *bicd-1* also failed to undergo pronuclear fusion (Fig S9). Moreover, *Drosophila D-TACC* and *C. elegans chc-1* and *tac-1* are also needed for pronuclear fusion (Fig S9). These genes might be required indirectly, through their role in meiosis, since preliminary data also suggest that meiosis II is compromised in *bicd-1* and *chc-1 ds*RNA fed worms (Fig S9). Alternatively, they might play a more direct role in pronuclear migration, which is known to depend on Dynein and MTs in bovine, primate, and *C. elegans* embryos (Gönczy et al., 1999; Payne et al., 2003). Figuring out their precise mechanistic involvement in pronuclear fusion is an interesting question for further studies.

## MATERIALS AND METHODS

### *Drosophila* stocks

*P{matα4-GAL-VP16}, Nos-Gal4:Vp16* ; *UAS-Clc::GFP, Histone::RFP* ; C(1;Y)^1^, y^1^w: y^+^/0 and *C(1)RM, y*^*1*^ *v*^*1*^*/0* stocks were obtained from the Bloomington *Drosophila* Stock Center (NIH P400D018537), Flies containing the genome rescue transgenes p*CHC3+* (Bazinet et al., 1993) and *4C-CHC* (Kasprowicz et al., 2008), were kindly provided by C. Bazinet and P. Verstreken, respectively. *d-tacc*^*1*^ and *Df 3R(110)*, covering the *tacc* locus, were provided by J. Raff (Gergely et al., 2000). *UAS-Chc::eGFP* and *UAS-Chc::mcherry* flies (Li et al., 2010) were provided by S. Bullock. Transgenic flies *pUASP-Chc-V5-K10-AttB*, and *pUASP-myc-chc-K10-AttB* were described previously (Vazquez-Pianzola et al., 2014). *Df(2L)Exel7068* (Exelixis) was used as *BicD* deficiency (BicD^*Df*^). The *BicD*^*null*^ allele, *BicD*^*r5*^, *Bic-D*^*mom*^, and *BicD::GFP* were described (Paré and Suter, 2000; Ran et al., 1994; Swan and Suter, 1996; Vazquez-Pianzola et al., 2014). *White (w)* flies were used as controls.

For the production of unfertilized eggs, virgin *w* females were crossed to *C(1;Y)*^*1*^, *y*^*1*^*w: y*^*+*^*/0* males to generate XO males that are phenotypically normal but sterile. XO males were then crossed to virgin females of the desired phenotype. Eggs laid by these females were not fertilized.

### DNA constructs and generation of transgenic flies

The generation of the flies expressing the *hb-deGRadFP* construct was as follows. We cloned the NSlmb-vhhGFP4 sequence (that comprises the F-box domain contained in the N-terminal region of the *Drosophila* supernumerary limbs (Slmb) protein fused to the GFP-binding nanobody VhhGFP4 sequence) described by Caussinus and colleagues (Caussinus et al., 2011) under the *hunchback* (*hb*) minimal maternal promoter containing the 5’-UTR leader of the maternal 3.2 Kb *hb* transcript and combined it with the *bcd* 3’-UTR, following the strategy used by Schulz and Tautz who used this to induce an artificial Hb gradient in embryos (Schulz and Tautz, 1995). The *Nslmb-vhhGFP4* sequence was PCR-amplified with specific primers bearing *BamHI* and *KpnI* sites using the plasmid pUAS-Nslmb-vhhGFP4 as the template (Caussinus et al., 2011).

Primer sequences were the following ones. Nano-GFP-sense-BamHI (GGATCCATGATGAAAATGGAGACTGACAAAAT) and Nano-GFP-anti-Kpn (GGTACCTTAGCTGGAGACGGTGACCTGGGTG). PCR products were subcloned into the same sites of the pCaSper-AttB vector to create the plasmid pCasper-AttB-Nslmb-vhhGFP4 (or pw+GFP nanobody). A 1.5 Kbp fragment of the *bicoid* 3’-UTR genomic region (Berleth et al., 1988) was amplified from a genomic library using specific primers containing KpnI and NotI restrictions sites. Primers were: bcd 3’UTR sense-Kpn I (GGTACCACGCGTAGAAAGTTAGGTCTAGTCC) and bcd 3’UTR anti-Not I (GCGGCCGCGCTAGTGCTGCCTGTACAGTGTCT). The insert was cloned by T-end ligation into the pCR2.1 TOPO vector and later subcloned into the KpnI/ NotI sites of pCasper-AttB-Nslmb-vhhGFP4 to generate the pCasper-AttB-Nslmb-vhhGFP4-bcd 3’-UTR construct (or pw+ GFP nanobody-bcd 3’-UTR). The minimal maternal *hunchback (hb)* promoter region together with the 5’-UTR leader of the maternal 3.2 Kb *hb* transcript were amplified using as template the Lac8.0 construct already described (Margolis et al., 1994) and kindly provided by Jim Posakony. Primers for this amplification contained flanking BglII and BamHI sites and were the following ones. Hb-pr-sense–BglII (AGATCTTCCGGATCAGCGGCGCTGATCCTGC), and Hb-pr-Anti-BamHI (GGATCCCTTGGCGGCTCTAGACGGCTTGCGGACAGTCCAAGTGCAATTC). Inserts were further cloned into the *BamHI* of Nslmb-vhhGFP4-bcd 3’-UTR to generate Hb-Nslmb-vhhGFP4-bcd 3’UTR (or pw+ Hb-GFP nanobody-bcd3’UTR). This final plasmid was injected into embryos containing the ZH-attP-14/3R-86F landing platform to generate flies expressing the *hb-deGradFP* construct.

The *Don Juan (Dj)::mCherry* construct to produce transgenic flies was generated as follows. mCherry region was PCR-amplified with primers containing BamHI and HindIII sites using pC4-SqhP-mCherry plasmid as the template (a gift from Romain Levayer). The reverse primer added a GS-rich region to use as a linker and the stop codon was removed. The amplified fragment was further subcloned into the BamHI/HindIII sites of the plasmid *pw+SNattB* (Koch et al., 2009) to generate the plasmid *pw+attB*-mCherry. The genomic region containing the minimal promoter, the 5’-UTR, and the ORF of *Dj* gene was PCR-amplified from genomic fly DNA using primers containing EcoRI and BamHI sites, and cloned into the same sites of *pw+attB*-mCherry to generate the final construct *pw+attB - Don Juan (Dj)::mCherry*. This construct was further injected into flies containing the ZH-attP-52 / 3L-64A landing platform for transgenesis.

### Hatching rate determination and embryo development

Hatching rates were scored as follows. Virgin females of the desired genotype were crossed to control *white* males. Females were allowed to lay eggs on agar plates for several hours or overnight. Around 100-200 embryos were marked in the plate and further developed for 48 h at 25°C. After 48 h, embryos that did not hatch were counted. For scoring embryo development, 30 min to 60 min or 2- to 5h old embryos were collected. Embryos were then fixed and stained to detect both α-tubulin and DNA to score the development stage.

### Western blots

The preparation of ovaries for Western blots was as follows. Seven pairs of ovaries were collected in 20 µl of SDS-sample buffer, boiled for 2 min., vortexed for 15 sec., boiled for another 8 min. Finally, they were loaded onto an SDS –PAGE. Western blots were performed using mouse monoclonal anti-BicD antibodies (a mixture of 1B11 and 4C2, 1:10 dilution, (Suter and Steward, 1991), mouse anti-α-tubulin (1:2,000 dilution of the cell culture’s supernatant, Developmental Studies Hybridoma Bank (DSHB)), rabbit anti-mammalian Chc (1:500 dilution, (Hirst et al., 2009)), rabbit anti-GFP (1:3,000 dilution, Immunokontact), rabbit anti-Clc (1:3,000, (Heerssen et al., 2008)), mouse monoclonal anti-Myc 9E10 (1:5 cell culture supernatant, DSHB), and rabbit anti D-TACC (1:10,000 dilution, (Kao and Megraw, 2009)). Primary antibodies were detected using horseradish peroxidase-conjugated secondary antibodies (GE Healthcare). To detect the deGRadFP expression from the *hb-deGRadFP* construct, anti-llama IgG-heavy and light chain antibodies (1:500 dilution, Bethyl) were used and developed using anti-goat IgG (H+L)-HRP conjugated antibodies (1:500 dilution, Invitrogen).

### Immunoprecipitations (IPs)

IPs were done essentially as previously described (Vazquez-Pianzola et al., 2011). 30 µl of protein-G Sepharose beads (Amersham) and 1 ml of embryo extracts were used per IP. Extracts were prepared from 0-20 h old embryos expressing a Chc::Myc fusion protein under a maternal Tubulin promoter. As a negative control, extracts were also prepared from wild-type embryos (OreR). For each IP, 1 ml of cell culture supernatant of monoclonal anti-BicD 1B11, anti-BicD 4C2, or anti-Myc 9E10 antibodies were bound to the beads. To IP TACC, a solution containing 2 µl of the polyclonal anti-TACC UT316 antibody (Kao and Megraw, 2009) diluted in 1 ml of PBS was used to bind to the protein-G beads. Beads were resuspended in 30 µl of sample buffer, and 7 to 15 µl per well was analyzed by Western blot.

### Immunostainings

Immunostainings of embryos were done using the following primary antibodies: mouse anti-BicD (a mix of 1B11 and 4C2, 1:10 dilution), mouse anti-Myc 9E10 (dilution 1:5, DSHB), mouse anti-Flag (1:200, Sigma), mouse anti-V5 (1:200, Invitrogen), rabbit anti-Cnn (1:500 (Heuer et al., 1995)), rabbit anti Clc (1:500 (Heerssen et al., 2008)), mouse monoclonal anti-α-tubulin DM1A (1:500, Sigma), rabbit anti α-tubulin (1:500, Abcam), rabbit anti-GFP (1:300, previously preabsorbed on embryos, Immunokontakt), rabbit anti D-TACC UT316 (1:1,000 (Kao and Megraw, 2009)), rabbit anti D-TACC (1:500 (Gergely et al., 2000)), rabbit anti-Msps (1:500 (Cullen et al., 1999)), mouse anti-Flag (1:200, Sigma, F3165), rabbit anti CID (1:400 (Buster et al., 2013)), rabbit anti p-Histone H3 (S10) (1:200, Cell Signalling), mouse anti p-Histone H3 (1:200, Cell Signalling), rabbit anti BubR1 (1:2,000 (Logarinho et al., 2004)), and rabbit anti Mad2 (1/1,000 (Logarinho et al., 2004)). To detect the hb-deGRadFP construct, anti-llama IgG-heavy and light chain (1:500 dilution, Bethyl) was developed anti-goat Alexa Fluor 680 (H+L) (Ivitrogen).

Secondary antibodies used were Cy3-conjugated goat anti-mouse, DyLight 647-conjugated goat anti-mouse or anti-rabbit antibodies (Jackson Immunoresearch), A488-conjugated goat anti-rabbit (Molecular Probes), Oregon Green 488 conjugated goat anti-mouse (Molecular Probes), donkey anti-mouse AF488 (Molecular Probes), AF568 conjugated donkey anti-mouse, and AF488 conjugated donkey anti-rabbit A488 (Molecular Probes). Nuclei were visualized by staining DNA with 2.5 µg/ml Hoechst 33258 (Molecular Probes) for 40 minutes during the final wash or incubated overnight when early meiosis in embryos was analyzed. Control immunostainings using only secondary antibodies were performed to detect unspecific binding of the secondary antibodies. For co-localization studies, control samples using only one of the primary antibodies and both secondary antibodies were performed to detect bleed through to the other channel. When detecting tagged proteins, samples of wild-type specimens were used as a control for unspecific binding of the anti-tag antibody. Embryos were either fixed with 4% paraformaldehyde (PFA) or with methanol, as indicated. Methanol fixation was mainly used to preserve the cytoskeleton structure and or reduce the cytoplasmic levels in BicD staining. Methanol fixation was done as previously described (Kellogg et al., 1988). When antigens were not well preserved in methanol fixations (for example, this was observed for the Chc fusion proteins), fixation with 4% paraformaldehyde (PFA) was used. To detect endogenous BicD::GFP in ovaries, ovaries were fixed with 4% PFA. For preserving the endogenous Clc::GFP signal, embryos were hand-devitellinized and fixed in a mixture of Heptane saturated with PFA for 15 min. For preserving the Dj:mcherry and Chc::mCherry signal, the embryos were fixed with MetOH. Stage 14 oocytes for immunostainings were prepared as previously described (Radford and McKim, 2016).

Images were analyzed with a Leica TCS-SP8 confocal microscope. Most of the spindle pictures represent Z-stack maximal intensity projections along the frames. MII and Anaphase II images were acquired such that the central aster signal was highest, but bellow saturation. Due to the strong/robust Clc signal in the cytoplasm, the spindle perpendicular position to the surface and the depth of the sample, it was not possible to analyze Clc localization at the different regions of the meiotic spindles by directly analyzing the Z-stack of maximal confocal projection images. The high cytoplasmic signal in the first planes masked the localization at the spindle in the deeper planes. Thus, to detect the presence of Clc along the tandem meiosis spindles, Z-stack images were processed to correct for depth and bleaching in Image J. All stacks were processed the same way. First, a crop area corresponding to the spindle of the same size was cropped in each image. Channels were then split and subjected to corrections separately. Each channel was smooth with a median filter radius of 1 to decrease the noise. The channel corresponding to the Clc staining had the most robust signal intensity loss through the sample’s depth. This channel was subject to bleach correction with a simple ratio fit to compensate for intensity attenuation in the image’s deeper stacks. Channels corresponding to α-tubulin and DNA staining did not lose so much intensity with the sample’s depth. They were subjected to an attenuation correction using an ImageJ plugin already described (Biot et al., 2008). After intensity correction, the final image was created by merging the channels previously subjected to maximal intensity Z-stack projection.

### Fluorescent *in situ* hybridization (FISH)

For mRNA FISH, RNA probes were prepared and hybridization experiments were performed as previously described (Vazquez-Pianzola et al., 2017). To detect deGRadFP mRNA, the *VhhGFP4* region of the *hb-deGRadFP* transgene was first amplified by PCR with primers containing T3 and T7 promoter sequences, and the plasmid pUAS-Nslmb-vhhGFP4 as a template (Caussinus et al., 2011). Primer sequences were the following ones. The sense primer containing the T3 promoter was: GGGGGGAATTAACCCTCACTAAAGGGAGAATGGATCAAGTCCAACTGGTGGAGT and the antisense primer containing the T7 promoter was: GGGGGGTAATACGACTCACTATAGGGAGATTAGCTGGAGACGGTGACCTGGGTG. Antisense and sense probes were generated using T7 and T3 RNA polymerase, respectively. A sense probe was used to detect unspecific background.

DNA FISH to *Drosophila* chromosomes was done mainly according to (Dernburg, 2000). DNA oligos hybridizing to repetitive regions of the Y (AATAC)_6_ and the 2^nd^ chromosome (AACAC)_7_ were ordered modified with Cy3 and Cy5 fluorophores at their 5’ end, respectively, from Microsynth. The sequences were already described (Dernburg, 2000). The sequence used to detect the X chromosome by hybridizing to TTT-TCC-AAA-TTT-CGG-TCA-TCA-AAT-AAT-CAT recognizes the 359-bp satellite block on the *D. melanogaster* X chromosome as well as minor variants on chromosome 3 was described by (Ferree and Barbash, 2009). A 5’ end Cy5 fluorophore-labeled probe was used. Embryos were fixed with methanol for DNA FISH. Probes were used at a final concentration of 5 ng/µl in hybridization buffer. The blocking buffer used (before adding the first antibody for detection of the desired proteins) was: 2x SSC, 0.5% BSA (molecular grade BSA from Biolabs or Acetylated BSA from Ambion), and Tween-20 at 0.1%. Donkey fluorescent-conjugated secondary antibodies were used as described in the immunofluorescence experiments.

### Yeast two-hybrid experiments

Two-hybrid plasmids containing the *Drosophila Egl* (Egl; CG4051) full-length (pEgl-AD and pEgl-BD), *BicD* full-length (pBicD-AD and pBicD-BD) and the *BicD* carboxy-terminal domain (CTD; amino acids 535–782) (pBicD(535–782)-BD and pBicD (535–782)-AD) were previously described (Rashpa et al., 2017). *D-TACC* (CG9765) full-length; and *Chc* (CG9012) full-length, a fragment containing amino acids 329-803, and another one bearing amino acids 329–542 were cloned into the pOAD and or pOBD2 vectors (Cagney et al., 2000) in-frame either with the activator domain (AD) or the DNA-binding domain (BD) sequence of GAL4, respectively, to create the “prey” plasmids, pD-TACC-AD, pChc-AD, pChc (329–803)-AD, and pChc(329–542)-AD, as well as the “bait” plasmids pChc-BD, pChc (329–803)-BD and pChc (329–542)-BD. To produce the constructs D-TACC-Ala and D-TACC-Asp, D-TACC Ser863 was mutated to an alanine and aspartic acid codon, respectively, by site-directed mutagenesis. Changes were verified by sequencing. Interactions between “bait” and “prey” proteins were detected following a yeast interaction-mating method using the strains PJ69-4a and PJ69-4alpha (Cagney et al., 2000).

For all cases, diploid cells containing both AD and BD constructs were selected in media lacking tryptophan and leucine (–(L,W), *Growth test*). To test for possible interactions, cells were further replica-plated in media lacking leucine, tryptophan and adenine (–(L, W, a), media lacking leucine, tryptophan, histidine, and adenine (-(L,W,H,a), or media lacking leucine, tryptophan, and histidine, and containing the indicated amounts of 3-Amino-1,2,4-triazole (3AT), (-(*Interaction test*). For each test, ten-fold serial dilutions are presented. Growth was scored after 4-5 days of growth at 30°C.

### *C. elegans* strains, RNAi and imaging

N2, JDU233 (*H2B::mCherry;α-tubulin::GFP*), VJ512, *chc-1(tm2866)III/hT2[bli-4(e937) let-?(q782) qIs48](I;III)*. Feeding of worms with dsRNA was performed on NGM plates containing 1mM IPTG and carbenicillin using clones from the Ahringer RNAi library (Kamath and Ahringer, 2003). For non-lethal phenotypes, the progeny was imaged after 60 hours of feeding of gravid adults at 20°C. For lethal phenotypes, L3/L4 larvae were transferred onto RNAi plates and incubated for 12 h at room temperature to obtain gravid adults. The empty vector L4440 was used as a control. Embryo dissections were performed as previously described (Bellanger et al., 2007), and embryos were imaged immediately on an agar pad using an AxioVision (Zeiss) microscope with a 100x NA 1.4 oil objective with a GFP and an mCherry filter set. One plane was acquired every 10 seconds using Visiview, and images were processed using ImageJ and Photoshop.

## ACKNOWLEDGEMENTS

We thank S. Bullock, C. Bazinet, J. Hirst, J. Raff, T. Melgraw, H. Ohkura, G. Rogers, C. Sunkel, M. Affolter, E. Caussinus, the Bloomington *Drosophila* Stock Center (NIH P400D018537), and the Developmental Studies Hybridoma Bank (created by the NICHD of the NIH and maintained at The University of Iowa, Department of Biology, Iowa City, IA 52242) for constructs, fly stocks and antibodies. We thank FlyBase (U41HG000739) for the *Drosophila* genomic resources. We thank Andrew Swan for his scientific advice on how to study female meiosis and Yury Belyaev for his help with image processing.

This work was supported by funding’s from the Swiss National Science Foundation (SNF, project grant 31003A_173188; www.snf.ch) and the University of Bern (www.unibe.ch) to BS, an SNF grant (project grant IZCOZ0_189884/31003A_176226) to P.M. and an Equal Opportunity grant from the Phil.-Nat. faculty to P.V.P. (www.unibe.ch).

## Author contributions

P.V.P., D. B., G.S., G. H., G. M., and D. H performed experiments. P.V.P., G. S., P.M., and G.H. analyzed data. P.V. P. and B.S. conceived the studies. G.S. and P. M. contributed to writing the results of the *C. elegans* experiments. P.V.P and B.S. wrote the manuscript.

